# Dynamic prioritisation of attention when external and internal demands compete

**DOI:** 10.64898/2026.08.06.743163

**Authors:** Daniela Gresch, Anna C. Nobre, Sage E.P. Boettcher, Melissa L.-H. Võ

## Abstract

Attention enables the prioritisation of task-relevant information, whether encountered in the external world or maintained as internal representations. While the mechanisms of attention have been extensively studied within perception and working memory, everyday cognition often confronts us with competing demands from both domains simultaneously. How attention is divided under such cross-domain demands remains poorly understood. Here, participants performed a combined perception and working-memory task in which an informative cue appeared between a working-memory display and a subsequent perceptual display. On most trials, this cue simultaneously indicated two items for prioritisation: either two encoded working-memory items or two upcoming perceptual items (within-domain attention), or one item from each domain (cross-domain attention). Combining behavioural measures with electroencephalography allowed us to characterise how dividing attention across domains shapes performance and neural activity relative to dividing attention within one domain. Behaviourally, we observed a striking asymmetry: perceptual performance benefited from cross-domain versus within-domain attention, whereas working-memory performance showed the opposite pattern. Mirroring this behavioural imbalance, time-resolved decoding revealed distinct cross-versus within-domain competition in both perception and working memory, suggesting that external and internal attention are neurally separable even when engaged simultaneously. This competition between domains was also evident in neural markers of spatial attentional orienting: while initially impaired during cross-domain attention, orienting progressively favoured external over internal information. Together, these findings show that when perception and working memory compete, attention is not divided evenly. Instead, cross-domain attention constitutes a distinct attentional state, with its own prioritisation dynamics and neural representation.

**Significance statement:** Perception and working memory are typically studied as separate targets of attention, yet real-world behaviour requires continuous negotiation between what is currently available in the external environment and what is held in mind. This study addresses how that negotiation unfolds when both sources of information become relevant at once. By directly comparing divided attention across versus within domains, we show that competition between perception and working memory is not neutral: it produces asymmetric behavioural consequences, distinct neural states, and a progressive bias in attentional prioritisation towards upcoming perceptual input. These findings suggest that cross-domain attention is not a balanced extension of within-domain attention, but a domain-biased prioritisation regime in which perception and working memory systematically trade off against each other.

## Introduction

Goal-directed behaviour requires attention to coordinate competing demands from the external environment and internal memory (Nobre & Gresch, 2025; van Ede et al., 2026; Verschooren et al., 2026). While external and internal attention have traditionally been studied in isolation, recent advances have moved towards a more comprehensive account, highlighting the importance of understanding the dynamic interplay between attentional domains. Specifically, emerging work has explored how attention shifts between external and internal domains as perceptual and mnemonic priorities unfold sequentially (Gresch, Boettcher, Gohil, et al., 2024; Gresch, Boettcher, van Ede, et al., 2024; Hammer et al., 2024; Verschooren et al., 2020; Wang et al., 2025). Yet, in many everyday situations, external and internal contents compete for priority concurrently – for example, when navigating dense traffic while keeping a recently seen road sign in memory. How attention is divided under such cross-domain competition remains poorly understood, limiting our understanding of how perception and memory interact when they place concurrent demands on attention.

Although direct evidence is limited, studies of divided attention within the external or internal domains provide useful guiding clues. Within the external domain, attention can be divided across multiple spatial locations simultaneously (Awh & Pashler, 2000; Cavanagh & Alvarez, 2005; Müller et al., 2003), however, this comes with a cost relative to focusing on a single location (Dowd & Golomb, 2019; Harrison et al., 2023). Similar findings have been reported within the internal domain: while dividing attention across a subset of multiple memory representations may provide some benefit relative to a no-prioritisation baseline, selectively focusing on a single memory item yields superior performance (DiPuma et al., 2023; Heuer & Schubö, 2016; Makovski & Jiang, 2007; Oberauer & Bialkova, 2009; Ueno & Allen, 2025).

Several factors may shape how perceptual and mnemonic information compete when both require attention concurrently. First, the extent of functional overlap between external and internal attention mechanisms may determine the strength of cross-domain competition: greater overlap should make competition across and within domains more comparable, whereas greater independence should weaken competition across domains. Second, attention may favour one domain at a given moment, prioritising its contents over those of the other. For example, external information may require prioritisation for immediate selection from the sensory stream, whereas internal contents are already integrated and separated from one another, allowing their prioritisation to remain flexible and reversible (Myers et al., 2018; van Moorselaar et al., 2015). Alternatively, attention may be inherently biased towards internal representations, since they maintain the relevant templates to guide future behaviour (Verschooren & Egner, 2023). Beyond processes directly driving competition, coordinating attention across sensory input and internal representations may place additional demands on control processes that are not required when attention is divided within one domain (cf. Nobre & Gresch, 2025).

To gain initial insight into cross-domain attention, we designed a task in which, on most trials, attention had to be directed simultaneously towards either two external inputs, two internal representations, or one of each. We complemented behavioural outcomes with electroencephalography (EEG) to examine the temporal dynamics of attention divided across versus within domains. Multivariate decoding analyses tracked neural representations, while lateralised posterior 8-12 Hz alpha activity indexed visuospatial orienting and the relative attentional priority of simultaneously relevant external and internal information.

We discovered that, relative to dividing attention within domains, cross-domain attention had asymmetric behavioural consequences, benefiting upcoming external information over previously encoded internal information. Consistent with this behavioural effect, the neural patterns associated with cross-domain attention differed from those observed when attention was divided within either perception or working memory, suggesting that external and internal attention remain neurally distinct even when simultaneously engaged. Lastly, although spatial attentional orienting was initially impaired during cross-domain attention, it progressively shifted towards external input. Together, these findings demonstrate that juggling concurrent attentional demands across perception and working memory differs from dividing attention within either domain at both behavioural and neural levels, with external information systematically prioritised over internal information.

## Methods

### Participants

The sample size was set to *n* = 25, based on previous studies with similar outcome measures (Gresch, Boettcher, Gohil, et al., 2024; Gresch, Boettcher, van Ede, et al., 2024; Gresch et al., 2025). Participants were aged between 18 to 39 years (mean age: 25; 17 female, 8 male), with 21 reporting being right-handed and four left-handed. Individuals provided written informed consent before participating in the study and were paid €15 per hour. The study was approved by the Psychology Ethics Committee at Ludwig-Maximilian University of Munich.

### Task and procedure

Participants performed a combined perception and working-memory task that required attention to be directed either towards a single external or internal item, divided between two items within one domain, or divided between two items across the domains (Figure 1A). On each trial, four randomly oriented bars were presented within four coloured circular placeholders. Two bars were presented early in the trial and had to be encoded into working memory, while the remaining two bars appeared later within the previously unoccupied placeholders and served as the upcoming perceptual input. Between the presentation of these internal working-memory and external perceptual displays, a centrally presented colour cue indicated the colour(s) of the placeholder(s) corresponding to the item(s) most likely to be tested at the end of the trial. This cue could indicate either one or two potentially relevant items. Single-item cues directed attention towards a single item, either in working memory (i.e., retro-cues) or in the upcoming perceptual display (i.e., pre-cues), and were 100% valid. In contrast, double cues directed attention towards either two perceptual items, two working-memory items, or one item from each domain, with one of the cued items subsequently probed for report. This design allowed us to compare reports of external and internal targets (target domain: external vs. internal) as a function of whether attention had been directed towards a single item, divided between two items within a domain, or divided across two items in different domains (cue type: single-item vs. (double-)within vs. (double-)cross).

**Figure 1.**
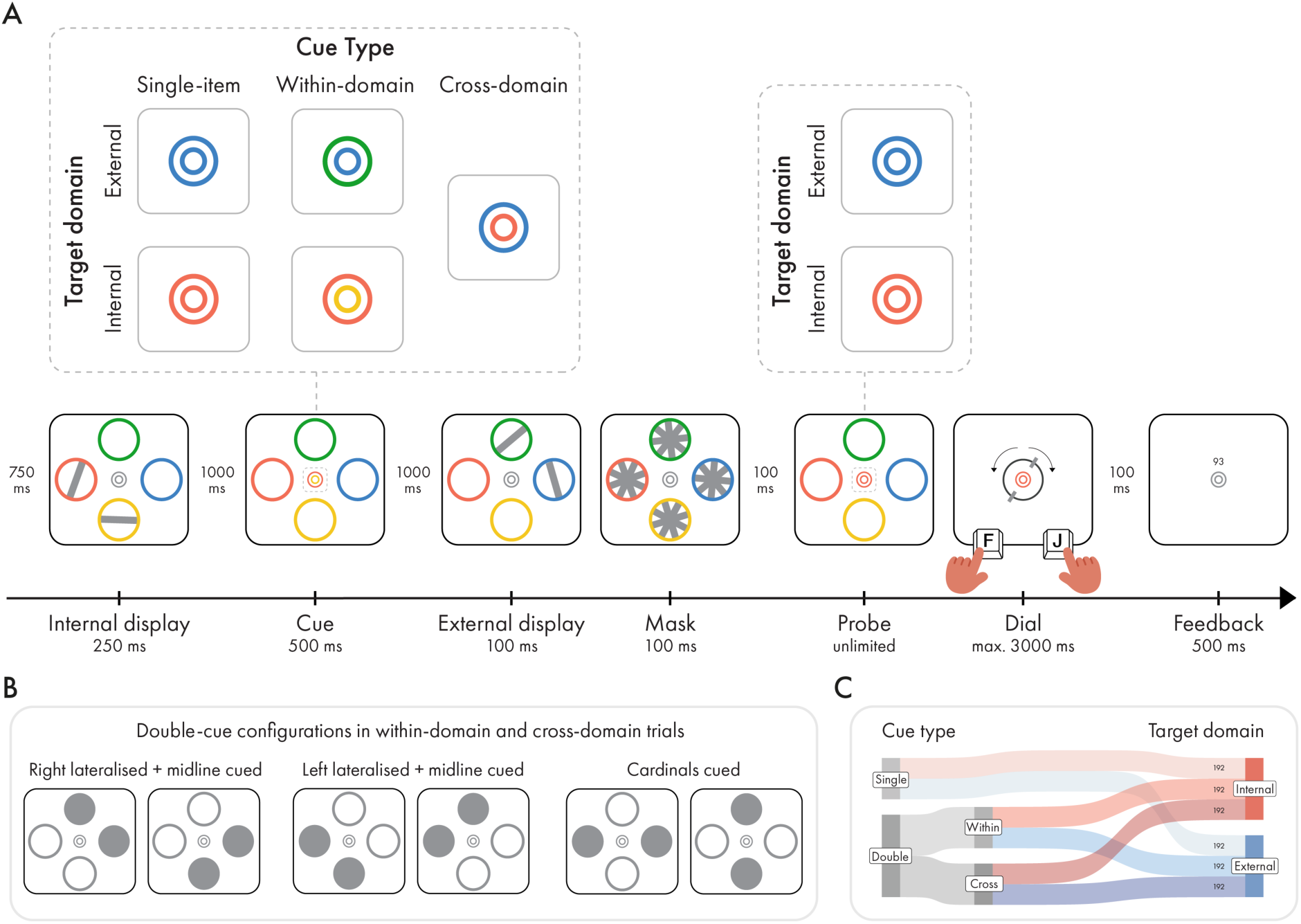
Task schematic and condition details. (A) Two tilted bars were presented early in the trial and encoded into working memory (internal display), whereas two further bars appeared later as perceptual input (external display). Between the internal and external displays, a centrally presented colour cue indicated the item(s) most likely to be tested. Single cues indicated one item in either the external or internal domain. Double cues indicated two potentially relevant items: either two items within the same domain (within-domain cues) or one external and one internal item (cross-domain cues). At the end of the trial, participants reported the tilt of the single cued item or, in double-cue trials, one of the two previously cued items. The content within the dashed rectangles shows enlarged views of the cue for clarity; the cue was presented at a smaller size in the experiment. (B) Double cues indicated one of six possible location configurations. (C) Ratio of conditions and number of trials across the experiment.

At the start of each trial, a grey central double-ring fixation marker (RGB value: [115, 115, 115]) was presented against a grey background (RGB value: [64, 64, 64]) for 750 ms. The inner and outer rings of the double-ring fixation marker had diameters of 0.25° and 0.5° of visual angle, respectively, and each ring had a width of 0.05° of visual angle. The double-ring fixation marker was surrounded by four coloured circular placeholders. Each of the placeholders appeared in one of four highly distinguishable colours (RGB values: blue [0, 159, 183], orange [218, 120, 0], green [120, 157, 0], pink [247, 35, 255]). Placeholders subtended 4° visual angle. They had a line width of 0.15° visual angle and were centred in the top, bottom, left, and right at a distance of 6.5° visual angle from fixation. The location mapping of the four placeholder colours was randomly determined on each trial. Placeholders stayed on the screen until the response was required at the end of the trial.

Next, two tilted grey bars (RGB value: [115, 115, 115]) subtending 0.5° × 3.89° appeared for 250 ms inside two of these placeholders (referred to as internal display). The orientation of the bars was randomly determined between 0 and 180°. Participants were instructed to memorize the orientation of these two bars. There were six possible configurations of the internal display, which occurred equally often throughout the experiment: the bars could appear in the placeholders located in the (1) top and left, (2) top and right, (3) top and bottom, (4) bottom and left, (5) bottom and right, and (6) left and right.

The internal display was followed by a 1000-ms delay, during which only the fixation marker and placeholders remained on the screen. The central fixation marker then changed colour for 500 ms to match the colour(s) of one or two placeholders, thereby acting as an informative cue indicating the item(s) relevant for the subsequent behavioural report.

In the *single-item-cue condition*, both the inner and outer rings of the fixation marker changed to the same colour. A single cue could indicate either a placeholder that had previously contained an item in the internal display (i.e., a retro-cue) or a placeholder that would contain an item in the upcoming external display (i.e., a pre-cue). Single cues were always 100% valid and therefore perfectly predicted the item that would be tested at the end of the trial.

In the *double-cue conditions*, the inner and outer rings of the fixation marker changed to different colours, each corresponding to a different placeholder. Three types of double cues were used. In the *internal within-domain condition*, both colours corresponded to placeholders that had contained items in the internal display. In the *external within-domain condition*, both colours corresponded to placeholders that would contain items in the upcoming external display. In the *cross-domain condition*, one colour corresponded to an item in memory and the other to an item in the upcoming external display. Independently, a double cue could indicate one of six possible placeholder combinations: (1) top and left, (2) top and right, (3) top and bottom, (4) bottom and left, (5) bottom and right, and (6) left and right (Figure 2B). The assignment of cue colours to the inner and outer rings was randomised. The test item was always selected from one of the two cued items, rendering the cue 100% valid at the level of the cued item set. For both single- and double-cue trials, cued locations and attentional domains were pseudo-randomised such that each placeholder location and each domain were cued equally often across the experiment.

**Figure 2.**
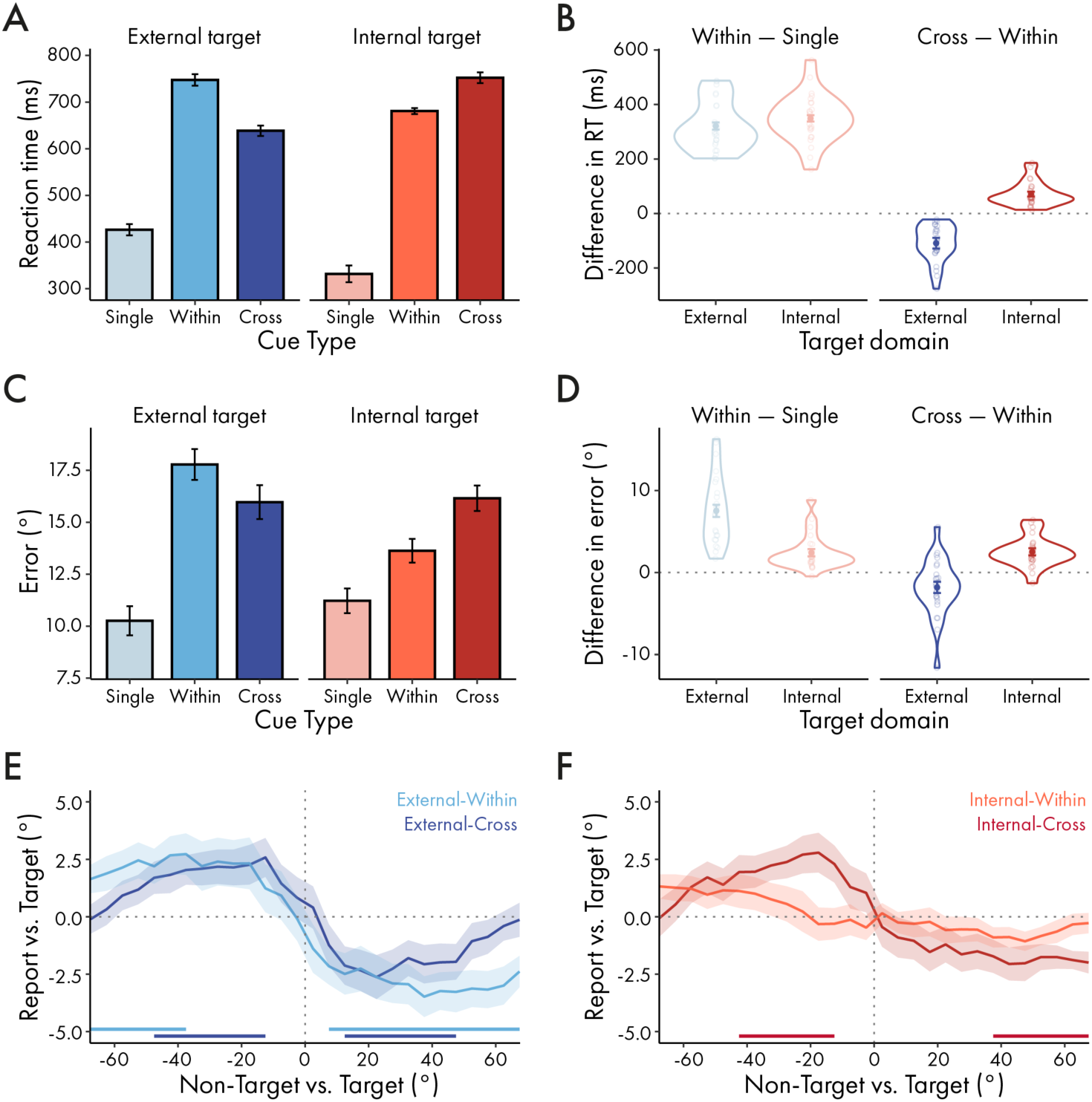
Behavioural performance. (A) Reaction times (RTs) as a function of cue type and target domain. (B) RT costs for within-domain versus single-item cues and cross-domain versus within-domain cues, shown separately for each target domain. (C) Same as (A), but for reproduction errors. (D) Same as (B), but for reproduction-error costs. (E) Average response bias as a function of the angular difference between the target and the cued non-target item in external-target trials. (F) Same as (E), but for internal-target trials. Error bars and shadings indicate *M* ± *SEM*. Horizontal lines indicate significant clusters.

The cue was followed by an inter-stimulus interval of 1000 ms before the onset of a perceptual display (referred to as external display), containing two randomly tilted bars. The new items appeared in the yet-unoccupied placeholders for 100 ms. For example, if the bars in the internal display appeared in the top and right placeholders, bars in the external display occurred the bottom and left placeholders. To make the perceptual discrimination challenging, items in the external array were masked. The masking array consisted of four overlayed titled bars presented in each of the placeholders for 100 ms. At each mask location, the four tilted bars differed 45° from the neighbouring bar. The overall orientation of each mask was randomly drawn at both 0 ms and 50 ms, thus creating the impression of a dynamic display. The masking parameters were set to equate the difficulty of external and internal reports, based on piloting efforts.

The offset of the dynamic mask was followed by a 100-ms delay, after which both the inner and outer rings of the fixation marker changed to the same colour (referred to as the probe), indicating which of the previously cued bars should be reported. Following probe onset, participants had unlimited time to initiate their response by pressing either the F or J key.

Once a response was initiated, the placeholders disappeared, and a visual response dial appeared around the fixation marker. The response dial matched the size of the placeholders and consisted of two markers positioned on opposite sides of a circle, corresponding to the ends of a bar. The dial always appeared in a vertical orientation and could be rotated leftwards or rightwards by pressing the left or right response key, respectively, at a rate of 0.1° per millisecond. Participants had 3000 ms to adjust the dial to match the remembered orientation of the target item. Once satisfied with their response, they pressed the space bar to confirm it.

After a 100-ms delay in which only the fixation marker stayed on the screen, participants received feedback in the form of a number ranging from 0 to 100, with 100 indicating a perfect report and 0 indicating that the adjusted orientation was perpendicular to the angle of the target item. If participants did not complete the orientation report within 3000 ms of response initiation, a "Too Slow" feedback message was presented. Feedback was presented 0.7° above the central fixation marker for 500 ms. Trials were separated by an inter-trial interval randomly drawn between 750 and 1000 ms. Between blocks, participants were presented with their average accuracy in the previous block.

The experiment consisted of 1152 trials completed across two sessions (Figure 1C). In each session, participants completed 16 blocks of 36 trials. One third of the trials (384) were single-cue trials, one third (384) were within-domain-cue trials, and one third (384) were cross-domain-cue trials. Within each cue condition, participants were equally likely to report an item from the internal display or the external display (192 trials each). Cue and target domains and locations, as well as the configurations of the external and internal displays, were counterbalanced across the experiment such that all permissible combinations occurred equally often, subject to the constraints imposed by the cue-target contingencies (e.g., if a within-domain cue indicated the top and bottom internal items, the target could only be selected from those two items). This yielded a total of 144 unique trial configurations. Each block contained an equal number of single-cue, within-domain-cue, and cross-domain-cue trials. For each cue condition within a block, six trials required participant to report an external item and six required them to report an internal item.

To familiarise themselves with the task, participants first completed practice blocks consisting of 12 trials, with equal numbers of each cue type and target domain. Practice continued until participants achieved an accuracy of at least 70%. Depending on performance, participants completed between two and eight practice blocks before reaching this criterion. The main experiment in each testing session took approximately 90 minutes to complete.

### Apparatus and data acquisition

Participants sat in front of a monitor (24-inch DELL U2412M; resolution 1920 × 1200 pixels; refresh rate 60 Hz; screen width 52 cm) at a viewing distance of 70 cm. An eye tracker (EyeLink 1000, SR Research) was positioned on the table approximately 15 cm in front of the monitor. Before acquisition, we calibrated the eye tracker using a 9-point calibration and validation method. Gaze was continuously tracked binocularly at a sampling rate of 1000 Hz, and participants were required to maintain fixation throughout the trial. The experimental script was generated using PsychoPy (version 2025.2.4; Peirce et al., 2019).

The EEG data were recorded using a 64-channel active electrode system (ActiCAP slim/snap & BrainAmp DC; Brain Products GmbH). Sixty-one electrodes were positioned across the scalp according to the international 10–10 system, with three additional electrodes placed below the left eye and at the left and right outer canthi to monitor vertical and horizontal eye movements via electrooculography (EOG). The reference electrode was placed on the left mastoid (i.e., behind the left ear) and the ground electrode was positioned on the forehead. During acquisition, data were low-pass filtered at 250 Hz and digitized at a sampling rate of 1000 Hz. Electrode impedances were at least below 10 kΩ and ideally below 5 kΩ. Data were referenced to the left mastoid during data acquisition.

### Behaviour: preprocessing

All data were preprocessed and analysed in the R statistical programming language (version 4.3.0; (R Core Team, 2025) using RStudio (version 2023.6.1.524; Posit team, 2022), following analysis procedures established in previous studies (Gresch et al., 2021, 2022, 2025). During preprocessing, trials were excluded if reaction times (RTs), measured from probe onset to the first key press, were more than 2.5 standard deviations (*SD*) above the participant’s mean RT across all conditions. Trials on which participants received a “Too Slow” feedback message during the orientation report were also excluded from further analyses. Participants for whom more than 10% of trials were excluded during preprocessing would have been removed from further analyses. No participant met this criterion. As an additional quality-control measure, datasets with a mean reproduction error of 40° or greater in any target domain × cue condition combination were flagged for potential exclusion. Reproduction error was defined as the mean absolute angular difference between the target and reported orientations. No dataset met this criterion. Following preprocessing, all 25 participants remained in the final sample. On average, 97.84% of trials (*SD* = 0.72) were retained per participant.

### Behaviour: response biases

We quantified whether the reported orientation exhibited a bias either towards (i.e., attractive bias) or away from (i.e., repulsive bias) the angle of the cued, non-target item. To achieve this, we first corrected for general response biases by subtracting each participant’s mean reproduction error from their error on individual trials. Next, reproduction errors were sorted and binned using a moving-window approach (step size = 5°, bin width = 45°) based on the orientation of the target relative to the cued non-target. The average response bias was calculated for each bin, resulting in a response-bias curve as a function of the target-to-non-target angular difference. This approach provided response-bias values indicating the degree of deviation of the report from the target orientation, with negative values reflecting an anticlockwise shift and positive values reflecting a clockwise shift. Similarly, the target-to-non-target angular difference was negative or positive depending on whether the non-target was oriented anticlockwise or clockwise relative to the target. An attractive bias occurred when the response-bias sign matched the target-to-non-target angular difference, while a repulsive bias was indicated when the signs were opposite.

### Behaviour: statistical analysis

The *lme4* package (Bates et al., 2015) was used to fit linear mixed-effects models (LMMs) using restricted maximum likelihood estimation (REML). To assess and improve the normality of the dependent variables, distributions were inspected, and candidate transformations were evaluated using the *MASS* (Venables & Ripley, 2002) and *bestNormalize* packages (Peterson, 2017). Based on repeated cross-validation, *bestNormalize* selected a Yeo-Johnson transformation for both response variables. The optimal transformation parameter was λ = 0.48 for RTs and λ = 0.05 for reproduction errors. Successive-difference (sliding) contrasts were specified for cue type (within-domain minus single-item; cross-domain minus within-domain) and target domain (internal minus external). Under this coding scheme, the intercept represents the grand mean, while the regression coefficients represent differences between adjacent factor levels in the specified direction.

Model selection began with the maximal random-effects structure justified by the design (Barr et al., 2013). Unless otherwise specified, this included by-subject random intercepts and random slopes for cue type, target domain, and their interaction. Potential overparameterisation was then assessed using a principal component analysis (PCA) of the random-effects variance-covariance matrix. Random slopes associated with unsupported dimensions identified by the PCA were removed if their exclusion did not significantly reduce model fit, as determined by likelihood-ratio tests (Bates et al., 2015). This approach resulted in a random-effects structure for RTs and reproduction errors that included a by-participant random intercept and maximal by-participant random slopes for cue type, target domain, and their interaction.

For LMMs, we report regression coefficients (β) with the *t*-statistic and *p*-values. *P*-values were estimated using Satterthwaite’s approximation degrees of freedom method as implemented in the *lmerTest* package (Kuznetsova et al., 2017). Statistical significance was assessed using a two-tailed α level of .05. Significant interactions were followed up with planned pairwise comparisons estimated using the *emmeans* package (Lenth & Piaskowski, 2017), with Bonferroni-adjusted *p*-values. Where applicable, differences in means between planned comparisons were compared using paired *t*-tests, and Cohen’s *d* was reported as an effect size measure.

This approach resulted in random-effects structures for both dependent variables (DVs; i.e., RTs and reproduction errors) that included by-participant random intercepts and maximal by-participant random slopes for cue type, target domain, and their interaction.

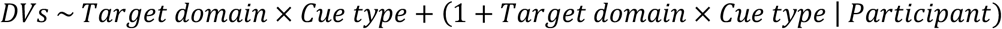

To statistically evaluate the response-bias analysis, we conducted a cluster-based permutation test with 10,000 iterations, comparing the response bias curve against zero.

### EEG: preprocessing

EEG data were analysed in Python using MNE-Python (version 1.11.0; Gramfort et al., 2013). The data were first notch-filtered at 50-Hz harmonics to remove line noise, band-pass filtered between 0.1 and 50 Hz, and downsampled to 250 Hz. The continuous data and power spectra were then visually inspected to identify noisy channels, which were removed prior to further preprocessing. On average, per session 0.74 ± 1.02 (*M* ± *SD*) electrodes per participant were identified as bad and subsequently interpolated. Next, EEG data were re-referenced to the common average. An independent component analysis (ICA) was used to identify and correct for artifacts associated with eye movements. For identifying ICA components associated with eye-movement artifacts, we correlated the time courses of all independent components with the measured horizontal and vertical EOG signal. Components with high correlation values and topographies characteristic of saccades or blinks were subsequently removed from the data. Data were then epoched from −200 to 1500 ms relative to the cue onset. No baseline correction was applied. We employed a generalized extreme studentized deviate (ESD) trial rejection approach to detect and exclude trials with high variance. On average, 26.22 ± 20.53 trials were removed per session, but never more than 10% of all trials. Finally, we applied a surface Laplacian transform (Perrin et al., 1989) to increase the spatial resolution of spectral modulations under consideration (as also done in: Boettcher et al., 2021; Gresch et al., 2025; Nasrawi et al., 2023; van Ede et al., 2019).

### EEG: decoding analysis

We performed time-resolved multivariate analysis on broadband EEG data using the Scikit-learn toolbox (version 1.8.0, Pedregosa et al., 2018) and built-in decoding functions provided by MNE-Python (version 1.11.0, Gramfort et al., 2013). The decoding analyses were implemented using a pipeline closely matching our previous approach (Gresch, Boettcher, Gohil, et al., 2024).

We extracted patterns of brain activity separately for each participant on a timepoint-by-timepoint basis. First, features (i.e., data at each time point for each electrode) were centred and scaled based on data from all trials. To reduce the number of redundant features for decoding, we performed Principal Component Analysis (PCA) on the standardised data while maintaining 99% of the variance. Training and testing were done on the same data using a 10-fold stratified cross-validation procedure. First, trials were randomized and divided into 10 equal-sized folds. Next, a leave-one-out procedure was used on the 10 folds, such that the classifier was trained on 9 folds and tested on the remaining fold. This procedure was repeated 10 times until each fold was used once for testing. A linear discriminant analysis (LDA) classifier was utilised to train on the provided training data and labels. LDA is a supervised dimensionality-reduction technique that aims to find a linear combination of features that maximizes class separability, and has been recommended for neuroscientific decoding studies (Grootswagers et al., 2017). We used the ROC-AUC as the scoring metric. This scoring metric takes into account the trade-off between true and false positive rates and is considered a sensitive, nonparametric, and criterion-free measure of classification (Hand & Till, 2001; Thölke et al., 2023). A ROC-AUC value of 0.5 means chance-level classification performance. The obtained classifier scores were averaged over all 10 folds, yielding a single decoding estimate per participant, timepoint, and condition. Lastly, decoding time courses were smoothed with a one-dimensional Gaussian filter with a *SD* of 10 samples (i.e., 40 ms).

To characterise the relationship between cue conditions, we additionally examined the classifier’s cross-validated prediction structure during the delay period, 500-1500 ms after cue onset. Using the same preprocessing and stratified 10-fold cross-validation procedure described above, we obtained held-out class predictions for each trial and used these to compute a condition-wise confusion matrix for each participant. Confusion matrices were row-normalised, such that each row reflected the proportion of trials from a given true condition classified as each predicted condition. Off-diagonal entries were then used to quantify the relative confusability between conditions. We computed a cross-domain-trial confusability index by subtracting cross-single confusability from cross-within confusability. Cross-within confusability was defined as the average of cross-domain trials classified as within-domain and within-domain trials classified as cross-domain, with cross-single confusability computed analogously. Positive values therefore indicate that cross-domain trials were more confusable with within-domain than with single-item trials.

In addition, we used temporal-generalization methods (King & Dehaene, 2014) to examine whether there are sustained neural activation patterns over time, that is, whether an activation pattern observed at one timepoint is observed again at a later timepoint. Generalization off the diagonal in the temporal generalization matrix is driven by overlap in the activation pattern at two different timepoints and therefore implies that the activation pattern is sustained in nature. Temporal generalization allows us to examine such potential overlaps in representations over time. This approach results in timepoint-by-timepoint decoding matrices in which each cell corresponds to a classification accuracy at a unique training- and test-timepoint combination.

To test which electrodes contributed most to the classifier likelihoods observed in our multivariate methods, we ran an EEG searchlight decoding analysis analogous to the searchlight analysis developed for functional MRI (Kriegeskorte et al., 2006). For this analysis, we applied the same decoding procedure described above to local clusters of four to eight neighbouring EEG electrodes (depending on electrode location), yielding a time-resolved decoding score for each electrode cluster. These decoding scores were averaged into time clusters of 150 ms each, resulting in ten topographical maps between 0 and 1500 ms. These topographical plots show whether decoding results were primarily driven by specific electrode clusters (Gresch, Boettcher, Gohil, et al., 2024; van Ede et al., 2019). We did not test any specific hypotheses regarding the spatial distributions of the effects and therefore did not run any statistical tests on the results.

### EEG: time-frequency analysis

We calculated time-frequency representations of power by convolving the data with Morlet wavelets from 3 to 40 Hz. The number of cycles increased linearly with frequency, corresponding to a constant wavelet duration of 400 ms and yielding an approximately constant temporal full-width at half-maximum (FWHM) of ∼150 ms and spectral FWHM of ∼5.9 Hz. Because the wavelets contained fewer than three cycles below ∼8 Hz, corresponding to a temporal FWHM of approximately one oscillatory cycle or less, estimates in this frequency range should be interpreted with caution (Cohen, 2019).

To obtain a time-resolved index of spatially selective changes in visual cortical excitability accompanying attentional orienting, we compared activity at posterior electrodes PO7/PO8 when the cued item was contralateral versus ipsilateral to each electrode. We expressed this contrast as a normalized difference (i.e., [(contra-ipsi)/(contra + ipsi)] × 100) and averaged it across the left and right electrodes. To derive time courses of lateralized visual activity, we averaged the contralateral versus ipsilateral response within the predefined alpha-frequency band (8-12 Hz). Topographies of lateralized visual activity were generated by contrasting trials in which cues indicated visual content in the right versus left hemifield, expressed as a normalized difference between right and left trials for each electrode (i.e., right minus left).

For the time-frequency analysis of contralateral versus ipsilateral power, we restricted the analysis to trials in which the cue indicated one lateralised item and one midline item (i.e., right + top, right + bottom, left + top, and left + bottom). This enabled a clear comparison of contralateral and ipsilateral activity, as alpha lateralisation most reliably indexes relative changes in visual excitability associated with shifts of spatial attention towards one hemifield and becomes ambiguous when both cued items occupy lateral positions.

It is important to note that restricting the analysis to trials in which one lateralised and one midline item were cued imposed specific constraints on the spatial configuration of the preceding internal display. In a within-domain trial that indicated two external items, one of the internal items was always present in the hemifield opposite to the cued external item. Thus, attention was consistently directed towards the hemifield opposite to the later-cued external item during encoding, resulting in pronounced alpha enhancement prior to cue onset. In contrast, on cross-domain trials (e.g., when a lateralised external item and a midline internal item were cued) the preceding internal display contained another item in the opposite hemifield on only half of the trials; on the remaining trials, the other internal item appeared at the midline. Attention was therefore directed towards the opposite hemifield during encoding on only half of the cross-domain trials, resulting in weaker lateralised pre-cue alpha activity than on within-domain trials, where attention was always directed towards that hemifield. Importantly, this was not an unintended design imbalance, as the decoding analyses, which included all trials, required cue and target domains, locations, and display configurations to be matched across conditions. Moreover, the pre-cue difference was unlikely to account for the alpha-lateralisation effect of interest, which emerged after cue-offset, allowing sufficient time for pre-existing activity to dissipate.

### EEG: statistical analysis

Statistical evaluation of EEG data used cluster-based non-parametric permutation, employing 10,000 permutations and a cluster alpha of 0.05. This approach sidesteps the problem of multiple comparisons in the statistical analysis of EEG data (Maris & Oostenveld, 2007). All statistical analyses were performed on unsmoothed data.

## Results

Participants performed a task in which attention was directed either towards a single external or internal item, divided between two items within the same domain, or divided across one external and one internal item across domains (Figure 1A). On each trial, participants first encoded two oriented bars into working memory. At the end of the trial, two additional bars were presented in placeholder locations. A colour cue appearing between the two displays indicated the item(s) most likely to be probed. Single-item cues indicated a singular external or internal item, whereas double cues indicated either two items from the same domain (within-domain cue) or one item from each domain (cross-domain cue). At the end of the trial, participants reproduced the orientation of the cued item in single-item trials or of one of the two cued items in within- and cross-domain trials. This design enabled comparisons of external and internal target reports as a function of attentional division.

### Dividing attention impairs performance in perception and working memory

We first examined the behavioural cost of dividing attention within a domain relative to focusing on a single item. Consistent with previous work, RTs were slower when two items within the same domain were cued than when only a single item was cued (β = 1.268, SE = 0.055, *t* = 22.850, *p* < 0.001; Figure 2A and 2B). This effect differed by target domain, as indicated by a significant cue type × target domain interaction (β = 0.277, SE = 0.050, *t* = 5.534, *p* < 0.001). Planned pairwise comparisons revealed significant differences in RTs between single-item and within-domain cue conditions for both external (β = 1.129, SE = 0.054, *z* = 21.086, *p* < 0.001) and internal target reports (β = 1.406, SE = 0.067, *z* = 20.868, *p* < 0.001). However, this RT difference was larger for internal than external target reports (*t*(24) = 5.531, *p* < 0.001, *d* = 1.106).

For reproduction errors, there was a robust effect of cue type, with larger errors when attention was divided within one domain than focused on a single item (β = 0.300, SE = 0.023, *t* = 13.022, *p* < 0.001; Figure 2C and 2D). This effect was qualified by a significant cue type × target domain interaction (β = −0.291, SE = 0.048, *t* = −6.016, *p* < 0.001). Planned pairwise comparisons showed that errors were larger following within-domain as compared to single-item cues for both external target reports (β = 0.446, SE = 0.039, *z* = 11.342, *p* < 0.001) and internal target reports (β = 0.155, SE = 0.026, *z* = 5.912, *p* < 0.001). However, the cost of dividing attention within a domain was significantly larger for external than internal reports, (*t*(24) = 6.010, *p* < 0.001, *d* = 1.202). Thus, our results replicate previous findings that, relative to focusing on a single item, dividing attention between items within either perception or working memory impairs performance. Going beyond previous work, we show that this cost takes different forms across domains, producing greater slowing for working-memory reports but greater accuracy costs for external reports. A complete summary of the inferential statistics is provided in Table S1 and S2.

### Cross-domain attention produces asymmetric trade-offs

Next, we turned to our main question: relative to dividing attention within a domain, does dividing attention across perception and working memory produce comparable consequences, impose additional cross-domain costs that impair both domains equally, or give rise to asymmetric trade-offs between external and internal reports?

Reaction times were overall faster when attention was divided across compared to within domains (β = −0.069, SE = 0.020, *t* = −3.535, *p* = 0.002; Figure 2A and 2B). Critically, this cross-within effect differed by target domain (β = 0.584, SE = 0.059, *t* = 9.933, *p* < 0.001). Specifically, planned pairwise comparisons revealed opposite effects for external and internal reports. For external reports, RTs were faster in cross-domain than within-domain trials (β = −0.361, SE = 0.043, *z* = −8.367, *p* < 0.001). In contrast, for internal reports, RTs were slower in within-domain than cross-domain trials (β = 0.223, SE = 0.025, *z* = 8.865, *p* < 0.001).

For reproduction errors, the effect of dividing attention across versus within domains also depended on target domain. (β = 0.273, SE = 0.049, *t* = 5.613, *p* < 0.001; Figure 2C and 2D). Again, planned pairwise comparisons showed opposite effects across target domains: for external reports, errors were larger in within-domain than cross-domain trials (β = −0.137, SE = 0.040, *z* = −3.416, *p* = 0.010), whereas for internal reports, errors were larger in cross-domain than within-domain trials (β = 0.137, SE = 0.024, *z* = 5.734, *p* < 0.001). Thus, dividing attention across domains neither produced consequences comparable to within-domain attention nor imposed a uniform additional cost. Instead, relative to the corresponding within-domain attention, cross-domain attention produced an asymmetric trade-off, improving external reports while impairing internal reports. A complete summary of the inferential statistics is provided in Table S1 and S2.

This pattern raises the possibility that externally cued non-target items exerted a stronger influence on report performance. Specifically, external information may have interfered with internal reports when attention was divided across domains, while competition between two external items may have produced larger costs when attention was divided within the external domain. To test whether the asymmetric error pattern reflected systematic distortions in report direction, we analysed reproduction biases, defined as biases in reports towards or away from the cued non-target item. Reproduction-bias analyses showed that external reports were repelled away from the other cued item in both within-domain (1^st^ cluster *p* = 0.022, 2^nd^ cluster *p* < 0.001) and cross-domain trials (1^st^ cluster *p* = 0.004, 2^nd^ cluster *p* = 0.017; Figure 2C). Internal reports, by contrast, were not biased in within-domain trials (*ps* > 0.125), but were repelled away from the external cued item in cross-domain trials (1^st^ cluster *p* = 0.018, 2^nd^ cluster *p* = 0.008; Figure 2F). However, report biases did not differ significantly between cross- and within-domain trials for either external (*ps* > 0.088) or internal reports (*ps* > 0.143), suggesting that directional response biases did not fully explain the asymmetric reproduction-error pattern.

### Cross-domain attention is neurally distinct from both external and internal within-domain attention

Behavioural analyses revealed asymmetric consequences of cross-domain attention: relative to within-domain attention, dividing attention across perception and working memory benefited external reports while impairing internal reports. These effects may reflect how attentional competition was resolved once the target was revealed at probe onset, rather than how attention was initially divided across domains. At cue onset, however, the target domain was still uncertain, meaning that all cross-domain trials imposed the same initial attentional demands regardless of whether perception or working memory was ultimately tested. We therefore used multivariate decoding of cue-locked neural activity to test whether dividing attention across versus within domains elicited distinct neural patterns during this initial period.

When contrasting cross-domain trials with a pooled within-domain class comprising external and internal within-domain trials, we observed distinct neural patterns for attention divided across versus within domains (cluster *p* < .001; Figure 3A). Since cross-domain trials prioritised only one item within each domain, they could conceivably have resembled single-item trials, which likewise involved prioritising a single item in one domain. The confusability analysis instead showed that cross-domain patterns were more similar to within-domain than to single-item patterns, *t*(24) = 11.870, *p* < .001, *d* = 2.374 (Figures 3B and 3C). Thus, despite being neurally distinguishable from within-domain attention, cross-domain attention more closely resembled a divided than a focused single-item attentional state.

**Figure 3.**
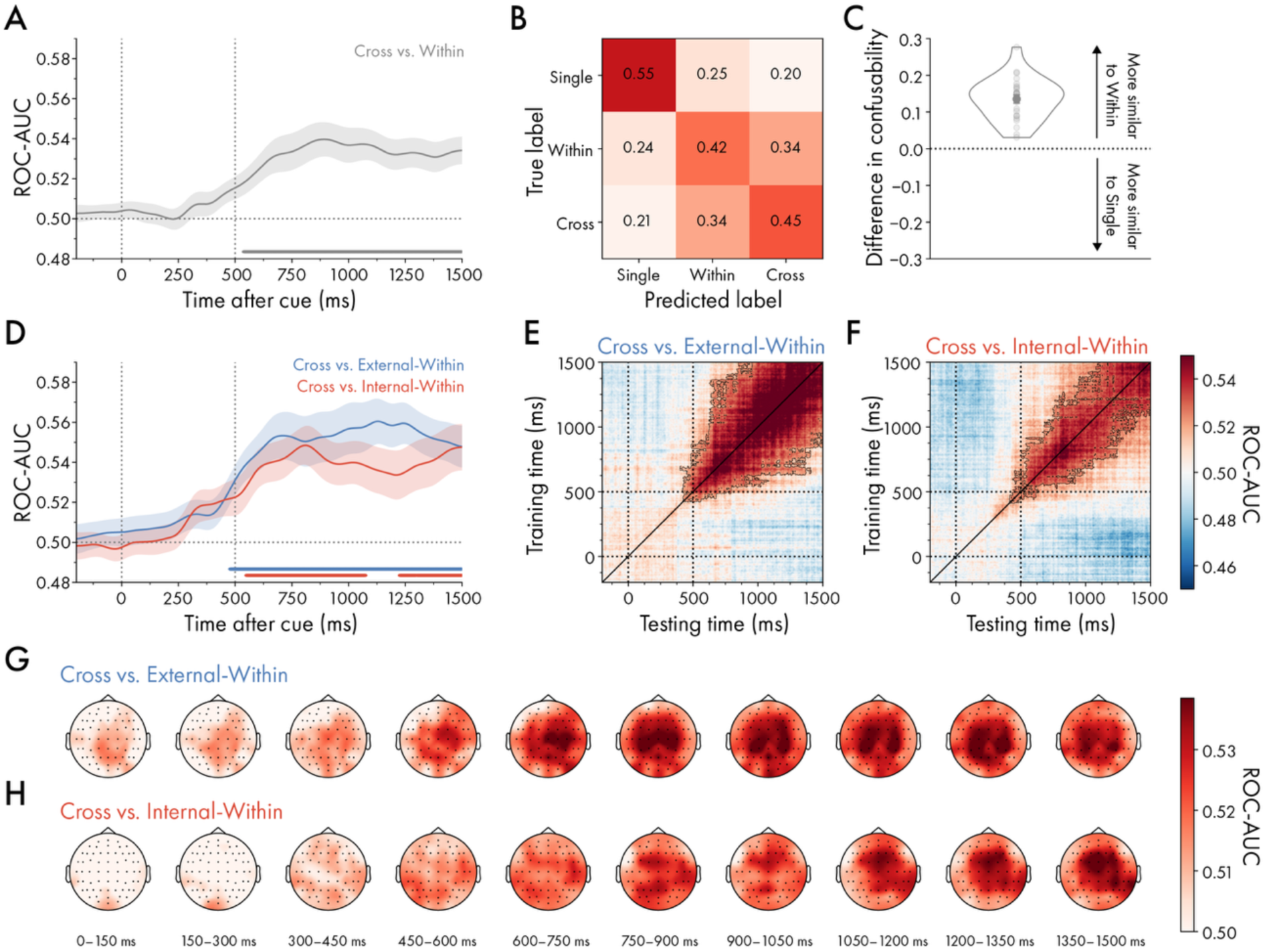
Time-resolved decoding of divided attention across versus within domains. (A) Average classifier performance for discriminating cross-from within-domain trials. (B) Confusion matrix showing classifier predictions across cue conditions from 500 to 1500 ms after cue onset. Rows indicate true conditions and columns indicate predicted conditions; off-diagonal values reflect classifier confusions between conditions. (C) Cross-domain confusability index computed as cross-within minus cross-single confusability. Positive values indicate greater confusability between cross-domain and within-domain than cross-domain and single-item trials. (D) Average classifier performance for discriminating cross-from external within-domain trials and cross-from internal within-domain trials. (E) Temporal generalisation matrix for discriminating cross-from external within-domain trials. The diagonal corresponds to the time course illustrated in (D). (F) Same as (E), but for cross-from internal within-domain trials. (G) Searchlight topographies averaged in 150-ms windows, showing the sensors contributing most strongly to cross-from external within-domain decoding. (H) Same as (G), but for cross-from internal within-domain decoding. Error bars and shadings indicate *M* ± *SEM*. Dots indicate individual participants. Cluster-permutation corrected significant time points are indicated with horizontal lines or black outlines.

Having established that cross-domain attention reflects a distinct yet divided attentional state, we next asked whether this difference holds when compared separately for external and internal within-domain attention. To test this possibility, we repeated the decoding analysis separately for each target domain. We trained and tested classifiers to distinguish cross-domain trials with a subsequent external target report from external within-domain trials, and cross-domain trials with a subsequent internal target report from internal within-domain trials. As the target item was not known at the time of the cue, this separation was performed retrospectively to equate trial numbers across analyses. Both classifiers showed reliable decoding (cross-domain vs. external within-domain: cluster *p* < 0.001; cross-domain vs. internal within-domain: 1^st^ cluster *p* < 0.001, 2^nd^ cluster *p* = 0.045, 3^rd^ cluster *p* < 0.001; Figure 3D), and decoding accuracy did not differ significantly between target domains (cluster *ps* > 0.469). Generalisation-across-time (GAT) analyses revealed reliable decoding for both cross-versus external within-domain (cluster *p* = 0.005) and cross-versus internal within-domain trials (cluster *p* = 0.018; Figure 3E and F), with temporal spread in both matrices suggesting some stability in the underlying neural patterns. However, no significant difference emerged in the GAT between cross-domain versus external within-domain and cross-domain versus internal within-domain trials (cluster *ps* > 0.636). Thus, decoding showed that cross-domain attention was neurally distinct from both divided external and divided internal attention, with no evidence that the pooled decoding was driven more strongly by either the external or internal within-domain condition.

Moreover, these findings extend previous evidence for distinct neural patterns of external and internal attention (Gresch, Boettcher, Gohil, et al., 2024), by showing that each remains distinguishable from a cross-domain attentional state in which both are engaged concurrently (see also Figure S1 for decoding of external and internal within-domain trials). Searchlight analyses further revealed qualitatively different scalp distributions for these distinctions (Figure 3G and 3H): decoding of cross-domain versus external within-domain trials was strongest over bilateral central-parietal channels, whereas decoding of cross-domain versus internal within-domain trials was more strongly expressed over frontocentral channels.

Together, these results demonstrate that dividing attention across domains produces distinct patterns of brain activity that differ from dividing attention within either the external or internal domain.

### Cross-domain attention initially impairs spatial orienting but ultimately favours external input

Broadband decoding showed that cross-domain attention differed from both external and internal within-domain attention. However, this analysis does not reveal how cross-domain attention prioritises information when external and internal representations compete directly. To address this, we examined lateralised 8-12 Hz alpha-band activity over posterior electrodes, using contra-versus ipsilateral differences as an index of spatial prioritisation in both the internal and external domain. We focused on double-cue trials in which one cued item was lateralised and the other appeared on the vertical midline. Specifically, in cross-domain trials, the cued external item occupied a left or right location while the cued internal item occupied a top or bottom location, or vice versa (Figure 1B). In within-domain trials, both cued items belonged to the same domain, with one positioned laterally and the other on the vertical midline. Combined with the analysis of alpha lateralisation, this cue configuration allowed us to isolate the prioritisation of external and internal separately while the two domains were in direct competition (also see Figure S2 for a comparison of within-domain and single-item trials).

Following the presentation of the simultaneous double cue, 8-12 Hz alpha-band activity showed clear lateralisation relative to the visual location of the lateralised cued item, with reduced contra-versus ipsilateral activity over PO7/PO8 in pooled within-domain and cross-domain trials (Figure 4A, 4D, and 4G). Interestingly, alpha lateralisation was initially stronger in within-domain than in cross-domain trials, suggesting an early attentional cost when attention was divided across domains (Figure 4J).

**Figure 4.**
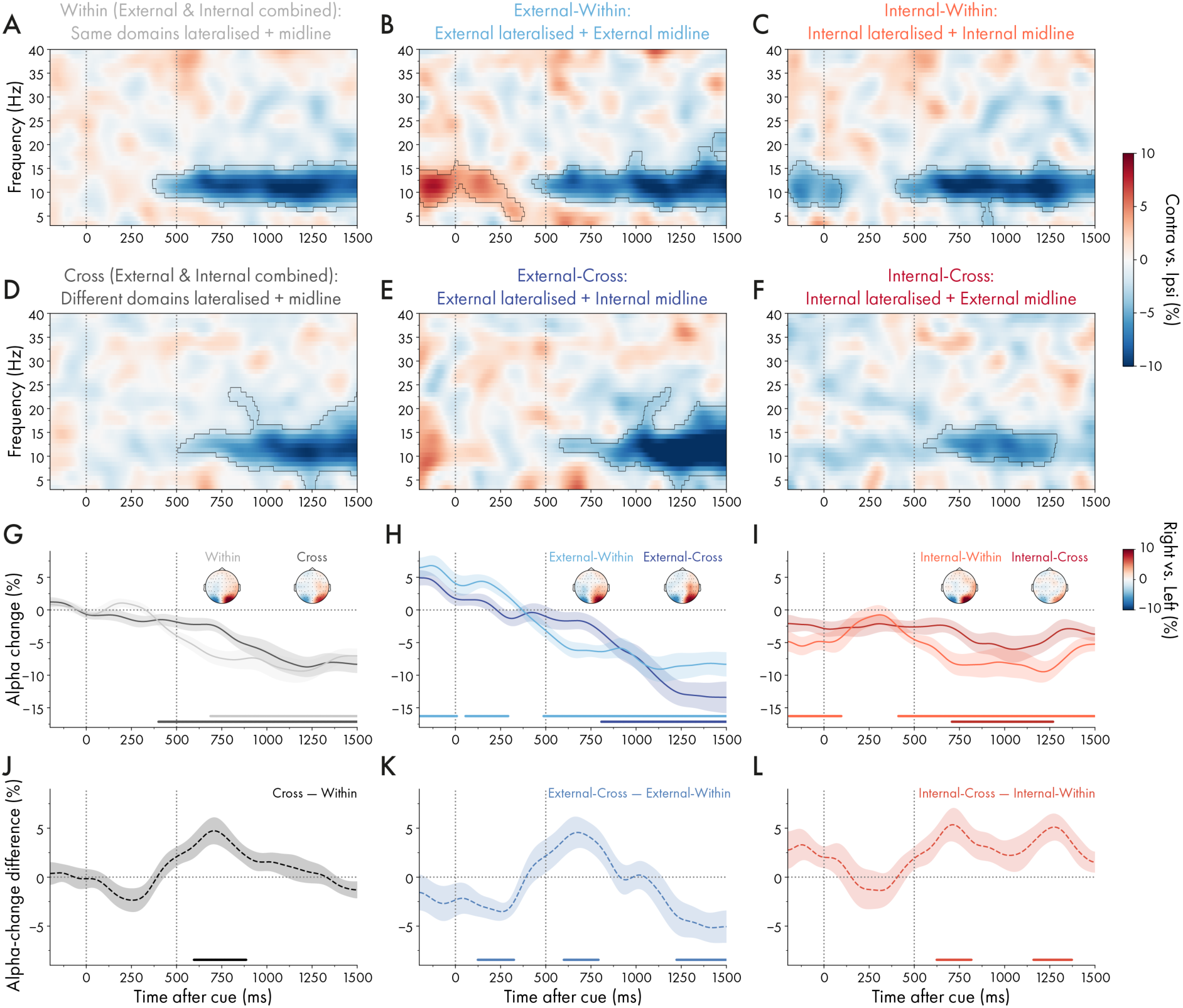
Lateralised neural activity following the cue in cross-domain and within-domain trials. (A) Lateralised time-frequency representation of contralateral versus ipsilateral activity relative to the location of the lateralised cued item, calculated over electrodes PO7/PO8. The analysis pooled within-domain trials in which one cued item was presented laterally and the other, from the same attentional domain, was presented on the midline. It included trials in which the lateralised item was either external or internal. (B) Same as (A), but for within-domain trials in which one cued external item was lateralised and the other cued external item was presented on the midline. (C) Same as (A), but for within-domain trials in which one cued internal item was lateralised and the other cued internal item was presented on the midline. (D) Same as (A but for pooled cross-domain trials in which one cued item was presented laterally and the other, from the opposite attentional domain, was presented on the midline. Trials with either an external or an internal lateralised item were included. (E) Same as (A), but for cross-domain trials in which the cued external item was lateralised and the cued internal item was presented on the midline. (F) Same as (A), for cross-domain trials in which the cued internal item was lateralised and the cued external item was presented on the midline. (G) Time courses of 8-12 Hz alpha-band lateralisation relative to the lateralised cued item in pooled within-domain and cross-domain trials. Topographies of the difference in 8-12 Hz alpha power between trials in which the cue indicated the selection of the right or left item, computed as right-minus-left cue trials and averaged across 500–1500 ms after cue onset. (H) Same as (G), but relative to the lateralised external cue in within-domain and cross-domain trials. (I) Same as (G), but relative to the lateralised internal cue in within-domain and cross-domain trials. (J) Difference between cross-domain and within-domain time courses pooled across trials in which either the external or internal cued item was lateralised. (K) Same as (J), but for trials in which the external cued item was lateralised. (L) Same as (J), but for trials in which the internal cued item was lateralised. Time courses and shadings indicate *M* ± *SEM*. Horizontal lines and black outlines indicate significant clusters.

When the domains were examined separately, this initial cross-domain vs. within-domain difference was present for both external (Figure 4B, 4E, 4H, and 4K) and internal attention (Figure 4C, 4F, 4I, and 4L), yet the pattern subsequently diverged between domains. For lateralised external items, alpha lateralisation became stronger in external-cross-domain than in external-within-domain trials towards the end of the delay. This indicates stronger prioritisation of the lateralised external item when it competed with an internal item on the vertical midline than when it competed with another external item. In contrast, for lateralised internal items, late alpha lateralisation was stronger in internal-within-domain than in internal-cross-domain trials, indicating weaker prioritisation when the lateralised internal item competed with a midline external item than when it competed with another internal item. These opposing effects suggest that prioritisation increasingly shifted towards the external location as the expected stimulus onset approached, accompanied by a corresponding reduction in the prioritisation of internal content.^1^

Together, these results reveal two temporally distinct consequences of cross-domain attentional competition. Initially, distributing attention across perception and working memory incurs a comparable cost to spatial prioritisation in both domains. Later, however, this pattern becomes asymmetric: upcoming perceptual information is prioritised more strongly under cross-domain than within-domain competition, whereas information held in working memory is deprioritised. A complete summary of the significant clusters is provided in Table S3.

## Discussion

Our study breaks new ground in understanding the mechanisms that govern attention under concurrent demands from perception and working memory. By independently manipulating whether attention was divided across or within domains and whether the final report concerned external or internal information, we were able to characterise both the behavioural consequences and the unfolding neural dynamics of cross-domain attention. Together, our findings reveal qualitatively different patterns of prioritisation and neural representation when attention is divided across as compared to within domains.

### Within-domain attention reveals differential costs across external and internal domains

Although the primary aim was to investigate cross-domain attention, our study additionally provided novel insight into the costs of dividing attention within individual domains. Within both perception and working memory, dividing attention impaired performance relative to focusing attention on a single item. This pattern is consistent with previous findings (Awh & Pashler, 2000; Cavanagh & Alvarez, 2005; DiPuma et al., 2023; Dowd & Golomb, 2019; Harrison et al., 2023; Heuer & Schubö, 2016; Makovski & Jiang, 2007; Oberauer & Bialkova, 2009; Ueno & Allen, 2025) and confirms that the double-cue manipulation successfully imposed a divided-attention cost. More importantly, our results extend these previous findings by showing that this cost manifests differently in perception and working memory. For internal reports, the cost was most pronounced in reaction times, suggesting that dividing attention across two working-memory representations particularly slowed access. For external reports, by contrast, the cost was most evident in reproduction errors, suggesting that dividing attention between two upcoming perceptual items more strongly reduced accuracy.

These effects may reflect fundamental differences in the nature of external and internal representations. Since working-memory representations have already been encoded, they may remain available with relatively high fidelity even when they are not initially prioritised. In line with this flexible and reversible nature of working memory, previous behavioural work has shown that initially uncued working-memory representations can still benefit from subsequent attentional prioritisation (Guo et al., 2024; Myers et al., 2018; Rerko & Oberauer, 2013; van Ede et al., 2017; van Moorselaar et al., 2015). The high fidelity of internal representations may be partly due to their orthogonalisation after encoding, which limits interference between them (Buschman, 2021; Panichello & Buschman, 2021; Piwek et al., 2023; Wan et al., 2022; Xu, 2024). The absence of systematic report biases between the two cued internal items accords with this interpretation. Moreover, if such orthogonalisation also reduces the need to actively maintain each representation in its original (spatial) format, prioritising a single item rather than dividing attention between two internal items may place similar demands on attention in working memory. This could also explain why alpha lateralisation did not differ between internal single-item and within-domain trials. By contrast, competing perceptual inputs have yet to be selected from the sensory stream. Dividing attention across multiple external inputs therefore limits the processing afforded to each, resulting in reduced perceptual precision (Carrasco, 2011).

### Cross-domain aLention reveals asymmetric prioritisation and distinct neural signatures

The primary aim of our study was to determine how attention is divided when perception and working memory impose concurrent demands. Our behavioural findings show a clear asymmetry: compared to within-domain attention, dividing attention across domains benefited external reports while impairing internal reports. One possible explanation for this finding is that cued external items generated stronger interference than cued internal items, regardless of the reported domain. Consistent with this possibility, the influence of perceptual input on working-memory representations has been shown to increase as the interval between memory encoding and subsequent perception lengthens beyond 100 ms (Teng et al., 2023). In our study, however, the report biases induced by the other cued non-target on double-cue trials did not directly mirror the asymmetric pattern observed in reproduction errors. Internal reports were repulsed away from the cued non-target item when it was external, but not when it was internal. By contrast, external reports were repulsed away from the cued non-target item regardless of its domain. Thus, cross-domain interference may have contributed to the observed asymmetry, at least for internal reports, but does not provide a complete explanation. The trade-off may therefore arise from additional mechanisms, including differences in the prioritisation of external and internal information.

Further insight into this possibility comes from the lateralisation of alpha-band activity at the time participants were prompted to divide attention. Towards the end of the delay interval, contra-versus ipsilateral alpha suppression was stronger for lateralised external items in cross-domain trials, where the other cued item was internal and midline, than in within-domain trials, where the other cued item was external and midline. Conversely, when the lateralised item was internal, contra-versus ipsilateral alpha suppression was weaker in cross-domain trials, where the other cued item was external and midline, than in within-domain trials, where the other cued item was internal and midline. Thus, attentional priority was progressively shifted towards upcoming perceptual information.

This pattern may reflect an adaptive strategy given the predictable temporal structure of the task. Temporal expectations allow attention to be dynamically directed towards external and internal information when each becomes most relevant (Dodwell et al., 2026; Echeverria-Altuna et al., 2024; van Ede et al., 2017; Williams et al., 2025). Since the external display was the next critical event following the cue, participants could enhance the selection of its contents by progressively shifting attentional priority towards external information while relying on the continued availability of the cued internal representation. Importantly, this external prioritisation was necessarily anticipatory: unlike internal cues, which could act directly on already encoded representations, external cues could only prepare attention for upcoming input but could not yet select a perceptual representation.

Additionally, the increased orienting of attention towards upcoming external information may arise because internal orienting is inherently more transient (Gresch, Boettcher, Gohil, et al., 2024; Myers et al., 2015; Wallis et al., 2015). This transient nature of internal attention may reflect the reformatting of internal representations after encoding – for example, into action-oriented codes (Boettcher et al., 2021; Myers et al., 2017; Rösner et al., 2022) – which may protect them from subsequent external interference (Gresch et al., 2021, 2025) and reduce the need for sustained spatial prioritisation.

Alpha-band lateralisation revealed not only a progressive shift towards external information over time, but also an initial cost of dividing attention across compared to within domains. Several complementary mechanisms may underlie this early cost. Dividing attention across domains may require translating the cue into separate control signals for perception and working memory, coordinating distinct representational formats, and/or aligning anticipatory orienting towards upcoming perceptual input with the prioritisation of an already encoded memory representation. The initial cost may therefore reflect a transient reconfiguration of attentional mechanisms across perceptual and mnemonic systems.

Finally, time-resolved decoding showed that different forms of divided attention were associated with unique patterns of brain activity. Cross-domain attention could be distinguished from the two within-domain conditions pooled together and from divided external and divided internal attention separately. Furthermore, the two within-domain conditions were also distinguishable from each other. These findings extend previous evidence that external and internal attention elicit distinct neural patterns when a single item is prioritised (Gresch, Boettcher, Gohil, et al., 2024), showing that this distinction persists even when attention is divided within each domain and that neither state resembles the concurrent engagement when divided across domains. This does not preclude external and internal attention from relying on partially overlapping neural mechanisms (Nobre et al., 2004; Zhou et al., 2022). Rather, the distinct decoding patterns may reflect differences both in the sensory and mnemonic representations on which attention operates and in the attentional operations these representations afford.

### Conclusion

In this study, we investigated how attention is divided across concurrent demands from perception and working memory. By directly comparing cross-domain with within-domain attention, we show that competition between perception and working memory is not neutral: it produces asymmetric behavioural consequences, distinguishable patterns of neural activity, and a progressive bias in spatial orienting towards upcoming perceptual input. Together, these findings reveal fundamentally different principles of prioritisation and neural representation when attention is divided across rather than within domains, opening new avenues for understanding the mechanisms through which attention bridges the external and internal worlds.

## Data and code availability

The experimental code, data, and analysis scripts are publicly available on the Open Science Framework (OSF): [will be shared upon acceptance].

## CRediT author contribution statement

Daniela Gresch: Conceptualization, Methodology, Software, Investigation, Formal analysis, Visualization, Writing – original draft, Funding acquisition. Anna C. Nobre: Conceptualization, Supervision, Writing – review & editing. Sage E.P. Boettcher: Supervision, Writing – review & editing. Melissa L.-H. Võ: Supervision, Writing – review & editing, Funding acquisition.

## Acknowledgements

This research was supported by an LMU Postdoc Support Fund awarded to D.G. Additional funding was provided by the Deutsche Forschungsgemeinschaft (DFG, German Research Foundation) under Germany’s Excellence Strategy EXC 3066 (“The Adaptive Mind”, Project No. 533717223) and through Research Unit FOR 5368 (“Abstract Representations in Neural Architectures”, Project No. 459426179), awarded to M.L.-H.V.

Moreover, the authors thank the research assistants at the Scene Grammar Lab – Johanna Gonan, Mercedes N. Hainzl Fernández, Vanessa Kauffmann, and Moritz Madysa – for their assistance with data collection.

## Supplementary Material

**Table S1.**
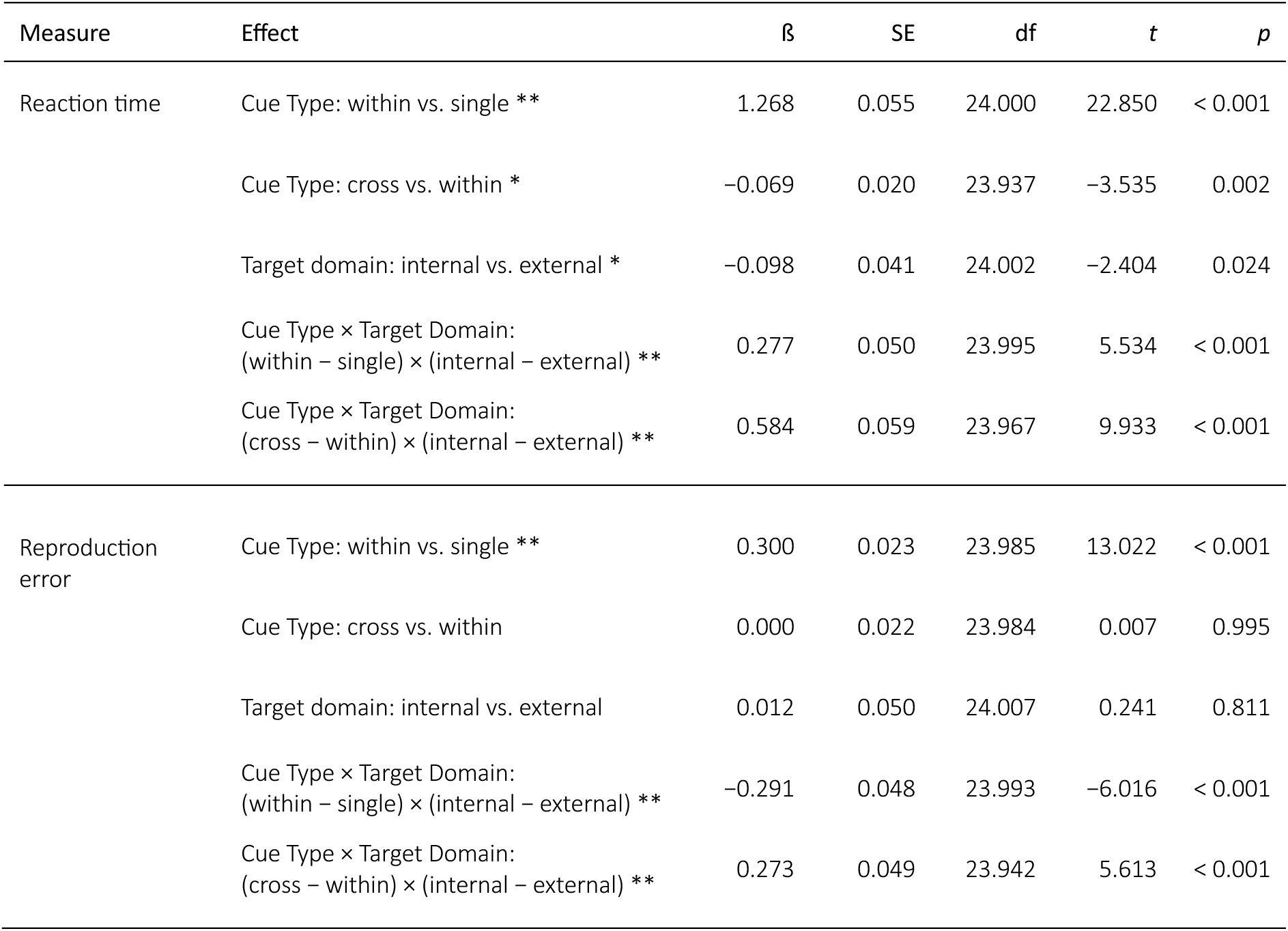
Outcomes of linear-mixed effects models (LMMs) analysing the effect of target domain and cue type on reaction times and reproduction errors. Contrasts were calculated as the first condition minus the second condition (within − single, cross − within, and internal − external). Interaction terms are differences-in-differences. * indicates p < 0.05, ** indicates p < 0.001.

| Measure | Effect | $\beta$ | SE | df | $t$ | $p$ |
| --- | --- | --- | --- | --- | --- | --- |
| Reaction time | Cue Type: within vs. single ** | 1.268 | 0.055 | 24.000 | 22.850 | < 0.001 |
|  | Cue Type: cross vs. within * | -0.069 | 0.020 | 23.937 | -3.535 | 0.002 |
|  | Target domain: internal vs. external * | -0.098 | 0.041 | 24.002 | -2.404 | 0.024 |
| | Cue Type × Target Domain:<br>(within – single) × (internal – external) ** | 0.277 | 0.050 | 23.995 | 5.534 | < 0.001 |
| | Cue Type × Target Domain:<br>(cross – within) × (internal – external) ** | 0.584 | 0.059 | 23.967 | 9.933 | < 0.001 |
| Reproduction error | Cue Type: within vs. single ** | 0.300 | 0.023 | 23.985 | 13.022 | < 0.001 |
|  | Cue Type: cross vs. within | 0.000 | 0.022 | 23.984 | 0.007 | 0.995 |
|  | Target domain: internal vs. external | 0.012 | 0.050 | 24.007 | 0.241 | 0.811 |
| | Cue Type × Target Domain:<br>(within – single) × (internal – external) ** | -0.291 | 0.048 | 23.993 | -6.016 | < 0.001 |
| | Cue Type × Target Domain:<br>(cross – within) × (internal – external) ** | 0.273 | 0.049 | 23.942 | 5.613 | < 0.001 |

**Table S2.** Pairwise comparisons between cue-type and target-domain conditions for reaction times and reproduction errors. Estimates and Cohen’s *d* were calculated as the first condition minus the second condition. Reported *p* values were adjusted for multiple comparisons using the Bonferroni correction. * indicates p < 0.05; ** indicates p < 0.001.

| Measure | Pairwise comparison | $\beta$ | SE | *z* | $p_{\text{Bonferroni}}$ |
| --- | --- | --- | --- | --- | --- |
| Reaction time | External-single vs. external-within ** | -1.129 | 0.054 | -21.086 | < 0.001 |
|  | External-single vs. external-cross ** | -0.768 | 0.051 | -14.955 | < 0.001 |
|  | External-single vs. internal-single** | 0.477 | 0.073 | 6.532 | < 0.001 |
|  | External-single vs. internal-within** | -0.929 | 0.051 | -18.091 | < 0.001 |
|  | External-single vs. internal-cross ** | -1.152 | 0.057 | -20.056 | < 0.001 |
|  | External-within vs. external-cross ** | 0.361 | 0.043 | 8.367 | < 0.001 |
|  | External-within vs. internal-single** | 1.606 | 0.096 | 16.656 | < 0.001 |
|  | External-within vs. internal-within** | 0.200 | 0.041 | 4.858 | < 0.001 |
|  | External-within vs. internal-cross | -0.023 | 0.037 | -0.614 | 1.000 |
|  | External-cross vs. internal-single** | 1.245 | 0.084 | 14.888 | < 0.001 |
|  | External-cross vs. internal-within* | -0.161 | 0.041 | -3.960 | 0.001 |
|  | External-cross vs. internal-cross ** | -0.384 | 0.048 | -7.994 | < 0.001 |
|  | Internal-single vs. internal-within** | -1.406 | 0.067 | -20.868 | < 0.001 |
|  | Internal-single vs. internal-cross ** | -1.629 | 0.085 | -19.211 | < 0.001 |
|  | Internal-within vs. internal-cross ** | -0.223 | 0.025 | -8.885 | < 0.001 |
| Reproduction error | External-single vs. external-within ** | -0.446 | 0.039 | -11.342 | < 0.001 |
|  | External-single vs. external-cross ** | -0.309 | 0.049 | -6.256 | < 0.001 |
|  | External-single vs. internal-single | -0.115 | 0.055 | -2.091 | .548 |
|  | External-single vs. internal-within** | -0.270 | 0.055 | -4.956 | < 0.001 |
|  | External-single vs. internal-cross ** | -0.407 | 0.059 | -6.869 | < 0.001 |
|  | External-within vs. external-cross * | 0.137 | 0.040 | 3.416 | 0.010 |
|  | External-within vs. internal-single** | 0.331 | 0.056 | 5.942 | < .001 |
|  | External-within vs. internal-within* | 0.176 | 0.056 | 3.133 | 0.026 |
|  | External-within vs. internal-cross | 0.039 | 0.057 | 0.691 | 1.000 |
|  | External-cross vs. internal-single* | 0.194 | 0.062 | 3.159 | 0.024 |
|  | External-cross vs. internal-within | 0.039 | 0.061 | 0.651 | 1.000 |
|  | External-cross vs. internal-cross | -0.097 | 0.063 | -1.557 | 1.000 |
|  | Internal-single vs. internal-within** | -0.155 | 0.026 | -5.912 | < 0.001 |
|  | Internal-single vs. internal-cross ** | -0.292 | 0.027 | -10.767 | < 0.001 |
|  | Internal-within vs. internal-cross ** | -0.137 | 0.024 | -5.734 | < 0.001 |

**Table S3.** Cluster-based permutation test results for alpha-band lateralisation. Results for the time-frequency representations correspond to Figures 4A-G, those for the time courses to Figures 4H-J, and those for the difference waves to Figures 4K-L.

|  | Combined |  | External |  | Internal |  |
| --- | --- | --- | --- | --- | --- | --- |
|  | Within | Cross | Within | Cross | Within | Cross |
| Time-frequency representations | $p < 0.001$ | $p < 0.001$ | $p < 0.001$<br>$p < 0.001$ | $p < 0.001$ | $p = 0.039$<br>$p < 0.001$ | $p < 0.001$ |
| Time courses | $p < 0.001$ | $p < 0.001$ | $p = 0.023$<br>$p = 0.023$<br>$p < 0.001$ | $p < 0.001$ | $p = 0.024$<br>$p < 0.001$ | $p < 0.001$ |
| Difference waves | $p < 0.001$ | | $p = 0.022$<br>$p = 0.048$<br>$p = 0.005$ | | $p = 0.041$<br>$p = 0.021$ | |

**Figure S1.**
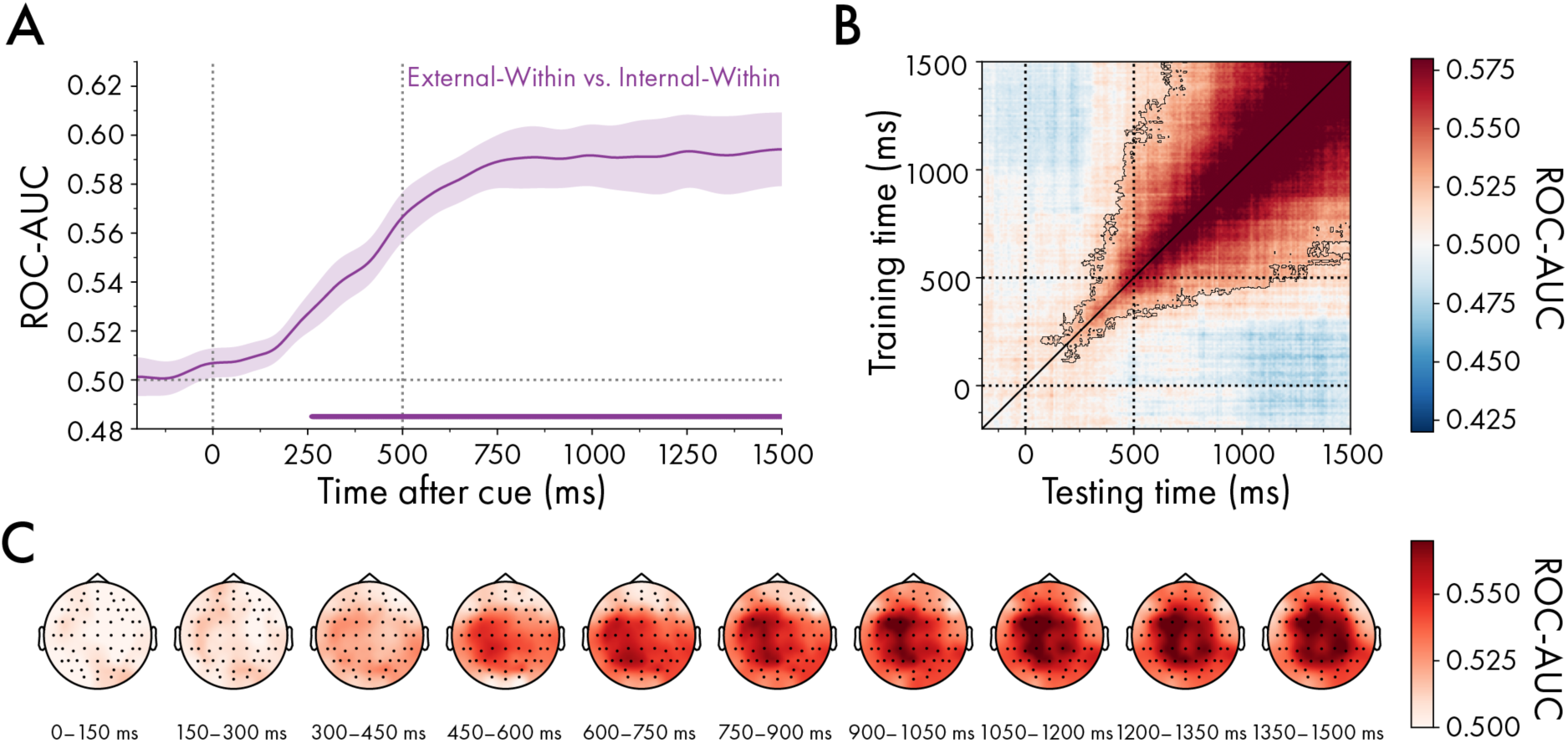
Time-resolved decoding of divided external and internal attention within domains. (A) Average classifier performance for discriminating external within-domain from internal within-domain trials (cluster *p* < 0.001). (B) Temporal generalisation matrix for discriminating external within-domain from internal within-domain trials (cluster *p* < 0.001). The diagonal corresponds to the time course illustrated in (A). (F) Same as (E), but for cross-from internal within-domain trials. (C) Searchlight topographies averaged in 150-ms windows, showing the sensors contributing most strongly to external within-domain from internal within-domain decoding. Error bars and shadings indicate *M* ± *SEM*. Cluster-permutation corrected significant time points are indicated with horizontal lines or black outlines.

**Figure S2.**
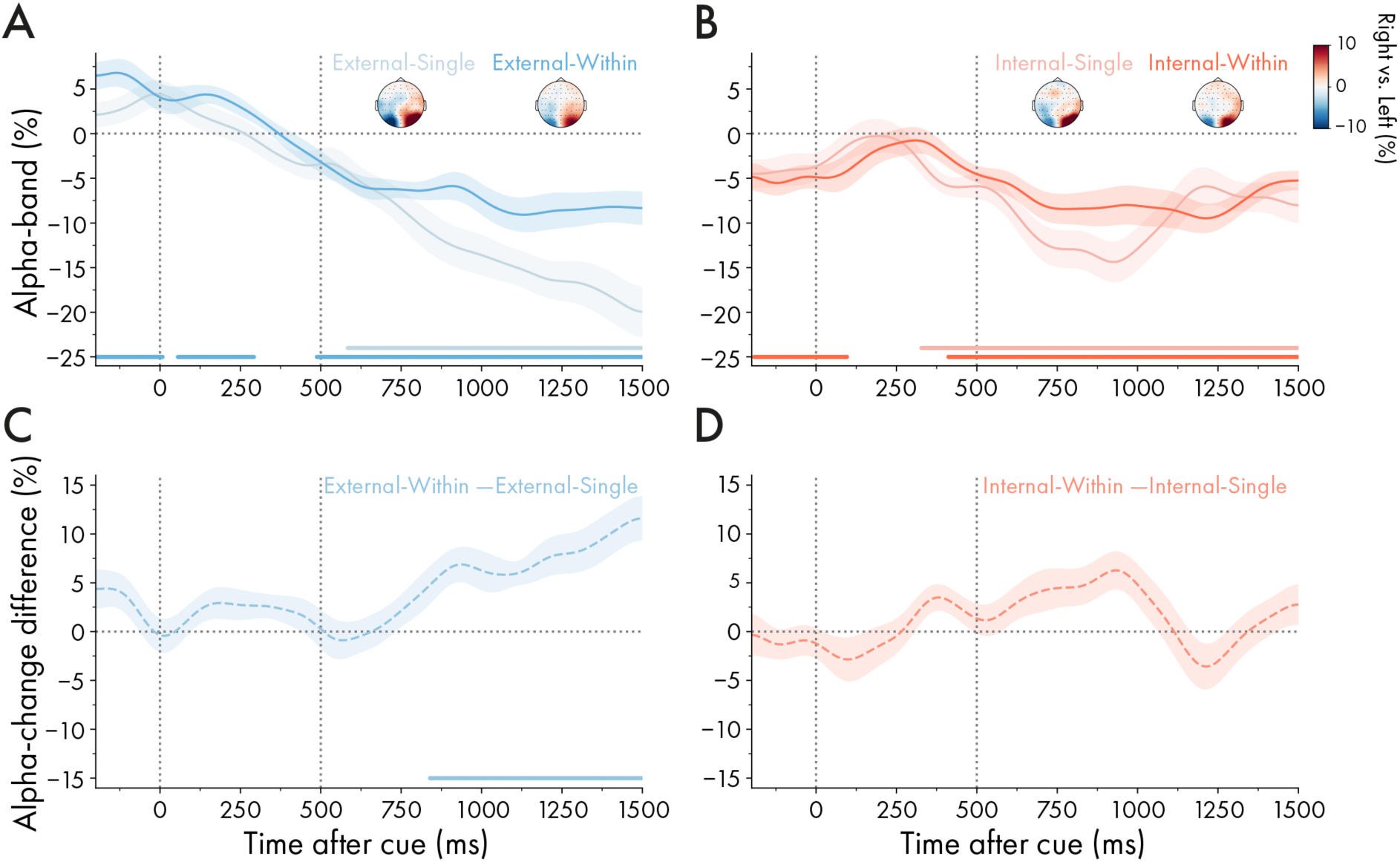
Lateralised neural activity following the cue in within-domain and single-item trials. (A) Time courses of 8-12 Hz alpha-band lateralisation relative to the lateralised cued external item in within-domain and single-item trials (single-item cluster: *p* < 0.001). Topographies of the difference in 8-12 Hz alpha power between trials in which the cue indicated the selection of the right or left item, computed as right-minus-left cue trials and averaged across 500–1500 ms after cue onset. (B) Same as (A), but relative to the lateralised cued internal item in within-domain and single-items trials (single-item cluster: *p* < 0.001). (C) Difference between within-domain and single-item time courses for trials in which the external cued item was lateralised (difference cluster: *p* < 0.001). (D) Same as (C), but for trials in which the internal cued item was lateralised (*p*s > 0.075). Time courses and shadings indicate *M* ± *SEM*. Horizontal lines and black outlines indicate significant clusters. See Table S3 for *p*-values of within-domain clusters.

## Footnotes

1 Additional clusters were observed prior to cue onset for external-within-domain and internal-within-domain trials. This effect reflected the display configurations across the trial rather than a confound in the cue-evoked lateralisation. In external-within-domain trials with one lateralised and one midline item, an internal item was always present in the hemifield opposite the later-cued external item, producing lateralised alpha enhancement relative to the subsequent external-item location. In internal-within-domain trials with one lateralised and one midline item, by contrast, the later-cued internal item had itself been encoded in the same hemifield, producing apparent lateralised alpha suppression prior to the cue onset relative to that item (see “Task and procedure” and “EEG: time-frequency analysis” for more information).

## References

Awh, E., & Pashler, H. (2000). Evidence for split attentional foci. Journal of Experimental Psychology. Human Perception and Performance, 26(2), 834–846. 10.1037//0096-1523.26.2.834

Barr, D. J., Levy, R., Scheepers, C., & Tily, H. J. (2013). Random effects structure for confirmatory hypothesis testing: Keep it maximal. Journal of Memory and Language, 68(3), 255–278. 10.1016/j.jml.2012.11.001

Bates, D., Mächler, M., Bolker, B., & Walker, S. (2015). Fitting Linear Mixed-Effects Models Using lme4. Journal of Statistical Software, 67, 1–48. 10.18637/jss.v067.i01

Boettcher, S. E. P., Gresch, D., Nobre, A. C., & van Ede, F. (2021). Output planning at the input stage in visual working memory. Science Advances, 7(13), eabe8212. 10.1126/sciadv.abe8212

Buschman, T. J. (2021). Balancing Flexibility and Interference in Working Memory. Annual Review of Vision Science, 7, 367–388. 10.1146/annurev-vision-100419-104831

Carrasco, M. (2011). Visual attention: The past 25 years. Vision Research, Vision Research 50th Anniversary Issue: Part 2, 51(13), 1484–1525. 10.1016/j.visres.2011.04.012

Cavanagh, P., & Alvarez, G. A. (2005). Tracking multiple targets with multifocal attention. Trends in Cognitive Sciences, 9(7), 349–354. 10.1016/j.tics.2005.05.009

Cohen, M. X. (2019). A better way to define and describe Morlet wavelets for time-frequency analysis. NeuroImage, 199, 81–86. 10.1016/j.neuroimage.2019.05.048

DiPuma, A., Lockhart, H. A., Emrich, S. M., & Ester, E. F. (2023). Retrospective cue benefits in visual working memory are limited to a single location at a time. ALention, Perception, & Psychophysics, 85(5), 1474–1485. 10.3758/s13414-023-02661-0

Dodwell, G., Nako, R., & Eimer, M. (2026). Rapid Changes of Attentional Priorities in Visual Search: Tracking Covert Switches of Preparatory Attentional Templates in Real Time. Journal of Cognitive Neuroscience, 38(5), 889–905. 10.1162/JOCN.a.2415

Dowd, E. W., & Golomb, J. D. (2019). Object-Feature Binding Survives Dynamic Shifts of Spatial Attention. Psychological Science, 30(3), 343–361. 10.1177/0956797618818481

Echeverria-Altuna, I., Boettcher, S. E. P., Ede, F. van, & Nobre, A. C. (2024). Dynamic prioritisation of sensory and motor contents in working memory (p. 2024.11.12.623203). bioRxiv. 10.1101/2024.11.12.623203

Gramfort, A., Luessi, M., Larson, E., Engemann, D., Strohmeier, D., Brodbeck, C., Goj, R., Jas, M., Brooks, T., Parkkonen, L., & Hämäläinen, M. (2013). MEG and EEG data analysis with MNE-Python. Frontiers in Neuroscience, 7. https://www.frontiersin.org/articles/10.3389/fnins.2013.00267

Gresch, D., Behnke, L., Ede, F. van, Nobre, A. C., & Boettcher, S. E. P. (2025). Neural dynamics of reselecting visual and motor contents in working memory after external interference. Journal of Neuroscience. 10.1523/JNEUROSCI.2347-24.2025

Gresch, D., Boettcher, S. E. P., Gohil, C., van Ede, F., & Nobre, A. C. (2024). Neural dynamics of shifting attention between perception and working-memory contents. Proceedings of the National Academy of Sciences, 121(47), e2406061121. 10.1073/pnas.2406061121

Gresch, D., Boettcher, S. E. P., Nobre, A. C., & van Ede, F. (2022). Consequences of predictable temporal structure in multi-task situations. Cognition, 225, 105156. 10.1016/j.cognition.2022.105156

Gresch, D., Boettcher, S. E. P., van Ede, F., & Nobre, A. C. (2021). Shielding working-memory representations from temporally predictable external interference. Cognition, 217, 104915. 10.1016/j.cognition.2021.104915

Gresch, D., Boettcher, S. E. P., van Ede, F., & Nobre, A. C. (2024). Shifting attention between perception and working memory. Cognition, 245, 105731. 10.1016/j.cognition.2024.105731

Grootswagers, T., Wardle, S. G., & Carlson, T. A. (2017). Decoding Dynamic Brain Patterns from Evoked Responses: A Tutorial on Multivariate Pattern Analysis Applied to Time Series Neuroimaging Data. Journal of Cognitive Neuroscience, 29(4), 677–697. 10.1162/jocn_a_01068

Guo, R., Wang, J., Fu, K., & Liu, Q. (2024). Exploring retro-cue effects on visual working memory: Insights from double-cue paradigm. Frontiers in Neuroscience, 17. https://www.frontiersin.org/articles/10.3389/fnins.2023.1338075

Hammer, J., Kajsova, M., Kalina, A., Krysl, D., Fabera, P., Kudr, M., Jezdik, P., Janca, R., Krsek, P., & Marusic, P. (2024). Antagonistic behavior of brain networks mediated by low-frequency oscillations: Electrophysiological dynamics during internal–external attention switching. Communications Biology, 7, 1105. 10.1038/s42003-024-06732-2

Hand, D. J., & Till, R. J. (2001). A Simple Generalisation of the Area Under the ROC Curve for Multiple Class Classification Problems. Machine Learning, 45(2), 171–186. 10.1023/A:1010920819831

Harrison, A. H., Ling, S., & Foster, J. J. (2023). The cost of divided attention for detection of simple visual features primarily reflects limits in post-perceptual processing. *ALention*, Perception & Psychophysics, 85(2), 377–386. 10.3758/s13414-022-02547-7

Heuer, A., & Schubö, A. (2016). Feature-based and spatial attentional selection in visual working memory. Memory & Cognition, 44(4), 621–632. 10.3758/s13421-015-0584-5

King, J.-R., & Dehaene, S. (2014). Characterizing the dynamics of mental representations: The temporal generalization method. Trends in Cognitive Sciences, 18(4), 203–210. 10.1016/j.tics.2014.01.002

Kriegeskorte, N., Goebel, R., & Bandettini, P. (2006). Information-based functional brain mapping. Proceedings of the National Academy of Sciences of the United States of America, 103(10), 3863–3868. 10.1073/pnas.0600244103

Kuznetsova, A., Brockhoff, P. B., & Christensen, R. H. B. (2017). lmerTest Package: Tests in Linear Mixed Effects Models. Journal of Statistical Software, 82, 1–26. 10.18637/jss.v082.i13

Lenth, R. V., & Piaskowski, J. (2017). emmeans: Estimated Marginal Means, aka Least-Squares Means (p. 2.0.3) [Dataset]. 10.32614/CRAN.package.emmeans

Makovski, T., & Jiang, Y. V. (2007). Distributing versus focusing attention in visual short-term memory. Psychonomic Bulletin & Review, 14(6), 1072–1078. 10.3758/BF03193093

Maris, E., & Oostenveld, R. (2007). Nonparametric statistical testing of EEG- and MEG-data. Journal of Neuroscience Methods, 164(1), 177–190. 10.1016/j.jneumeth.2007.03.024

Müller, M. M., Malinowski, P., Gruber, T., & Hillyard, S. A. (2003). Sustained division of the attentional spotlight. Nature, 424(6946), 309–312. 10.1038/nature01812

Myers, N. E., Chekroud, S. R., Stokes, M. G., & Nobre, A. C. (2018). Benefits of flexible prioritization in working memory can arise without costs. Journal of Experimental Psychology. Human Perception and Performance, 44(3), 398–411. 10.1037/xhp0000449

Myers, N. E., Stokes, M. G., & Nobre, A. C. (2017). Prioritizing Information during Working Memory: Beyond Sustained Internal Attention. Trends in Cognitive Sciences, 21(6), 449–461. 10.1016/j.tics.2017.03.010

Myers, N. E., Walther, L., Wallis, G., Stokes, M. G., & Nobre, A. C. (2015). Temporal dynamics of attention during encoding versus maintenance of working memory: Complementary views from event-related potentials and alpha-band oscillations. Journal of Cognitive Neuroscience, 27(3), 492–508. 10.1162/jocn_a_00727

Nasrawi, R., Boettcher, S. E. P., & van Ede, F. (2023). Prospection of Potential Actions during Visual Working Memory Starts Early, Is Flexible, and Predicts Behavior. Journal of Neuroscience, 43(49), 8515–8524. 10.1523/JNEUROSCI.0709-23.2023

Nobre, A. C., Coull, J. T., Maquet, P., Frith, C. D., Vandenberghe, R., & Mesulam, M. M. (2004). Orienting Attention to Locations in Perceptual Versus Mental Representations. Journal of Cognitive Neuroscience, 16(3), 363–373. 10.1162/089892904322926700

Nobre, A. C., & Gresch, D. (2025). How the brain shifts between external and internal attention. Neuron, 113(15), 2382–2398. 10.1016/j.neuron.2025.06.013

Oberauer, K., & Bialkova, S. (2009). Accessing information in working memory: Can the focus of attention grasp two elements at the same time? Journal of Experimental Psychology. General, 138(1), 64–87. 10.1037/a0014738

Panichello, M. F., & Buschman, T. J. (2021). Shared mechanisms underlie the control of working memory and attention. Nature, 592(7855), 601–605. 10.1038/s41586-021-03390-w

Pedregosa, F., Varoquaux, G., Gramfort, A., Michel, V., Thirion, B., Grisel, O., Blondel, M., Müller, A., Nothman, J., Louppe, G., Prettenhofer, P., Weiss, R., Dubourg, V., Vanderplas, J., Passos, A., Cournapeau, D., Brucher, M., Perrot, M., & Duchesnay, É. (2018). *Scikit-learn: Machine Learning in Python* (arXiv:1201.0490). arXiv. 10.48550/arXiv.1201.0490

Peirce, J., Gray, J. R., Simpson, S., MacAskill, M., Höchenberger, R., Sogo, H., Kastman, E., & Lindeløv, J. K. (2019). PsychoPy2: Experiments in behavior made easy. Behavior Research Methods, 51(1), 195–203. 10.3758/s13428-018-01193-y

Perrin, F., Pernier, J., Bertrand, O., & Echallier, J. F. (1989). Spherical splines for scalp potential and current density mapping. Electroencephalography and Clinical Neurophysiology, 72(2), 184–187. 10.1016/0013-4694(89)90180-6

Peterson, R. A. (2017). bestNormalize: Normalizing Transformation Functions (p. 1.9.2) [Dataset]. 10.32614/CRAN.package.bestNormalize

Piwek, E. P., Stokes, M. G., & Summerfield, C. (2023). A recurrent neural network model of prefrontal brain activity during a working memory task. PLOS Computational Biology, 19(10), e1011555. 10.1371/journal.pcbi.1011555

Posit team. (2022). RStudio: Integrated Development Environment for R. Posit Software, PBC. http://www.posit.co/

R Core Team. (2025). *R: A Language and Environment for Statistical Computing* [Dataset]. R Foundation for Statistical Computing.

Rerko, L., & Oberauer, K. (2013). Focused, unfocused, and defocused information in working memory. *Journal of Experimental Psychology: Learning*, Memory, and Cognition, 39, 1075–1096. 10.1037/a0031172

Rösner, M., Sabo, M., Klatt, L.-I., Wascher, E., & Schneider, D. (2022). Preparing for the unknown: How working memory provides a link between perception and anticipated action. NeuroImage, 260, 119466. 10.1016/j.neuroimage.2022.119466

Teng, C., Kaplan, S. M., Shomstein, S., & Kravitz, D. J. (2023). Assessing the interaction between working memory and perception through time. *ALention, Perception*, & Psychophysics. 10.3758/s13414-023-02785-3

Thölke, P., Mantilla-Ramos, Y.-J., Abdelhedi, H., Maschke, C., Dehgan, A., Harel, Y., Kemtur, A., Mekki Berrada, L., Sahraoui, M., Young, T., Bellemare Pépin, A., El Khantour, C., Landry, M., Pascarella, A., Hadid, V., Combrisson, E., O’Byrne, J., & Jerbi, K. (2023). Class imbalance should not throw you off balance: Choosing the right classifiers and performance metrics for brain decoding with imbalanced data. NeuroImage, 277, 120253. 10.1016/j.neuroimage.2023.120253

Ueno, T., & Allen, R. J. (2025). Running after two hares in visual working memory: Exploring retrospective attention to multiple items using simulation, behavioral outcomes, and eye tracking. Journal of Experimental Psychology. Human Perception and Performance, 51(3), 405–420. 10.1037/xhp0001270

van Ede, F., Chekroud, S. R., Stokes, M. G., & Nobre, A. C. (2019). Concurrent visual and motor selection during visual working memory guided action. Nature Neuroscience, 22(3), Article 3. 10.1038/s41593-018-0335-6

van Ede, F., Gresch, D., & Nobre, A. C. (2026). Disambiguating dimensions of external and internal brain processes. Trends in Neurosciences, 49(1), 5–7. 10.1016/j.tins.2025.10.001

van Ede, F., Niklaus, M., & Nobre, A. C. (2017). Temporal Expectations Guide Dynamic Prioritization in Visual Working Memory through Attenuated α Oscillations. The Journal of Neuroscience, 37(2), 437–445. 10.1523/JNEUROSCI.2272-16.2016

van Moorselaar, D., Olivers, C. N. L., Theeuwes, J., Lamme, V. A. F., & Sligte, I. G. (2015). Forgotten but not gone: Retro-cue costs and benefits in a double-cueing paradigm suggest multiple states in visual short-term memory. *Journal of Experimental Psychology: Learning*, Memory, and Cognition, 41, 1755–1763. 10.1037/xlm0000124

Venables, W. N., & Ripley, B. D. (2002). Modern Applied Statistics with S.

Verschooren, S., Dahl, M. J., Aly, M., & Mittner, M. (2026). Transition dynamics of external and internal attention across on-task and off-task states. Nature Reviews Psychology, 1–17. 10.1038/s44159-026-00578-7

Verschooren, S., & Egner, T. (2023). When the mind’s eye prevails: The Internal Dominance over External Attention (IDEA) hypothesis. Psychonomic Bulletin & Review. 10.3758/s13423-023-02272-8

Verschooren, S., Pourtois, G., & Egner, T. (2020). More efficient shielding for internal than external attention? Evidence from asymmetrical switch costs. Journal of Experimental Psychology: Human Perception and Performance, 46(9), 912–925. 10.1037/xhp0000758

Wallis, G., Stokes, M., Cousijn, H., Woolrich, M., & Nobre, A. C. (2015). Frontoparietal and Cingulo-opercular Networks Play Dissociable Roles in Control of Working Memory. Journal of Cognitive Neuroscience, 27(10), 2019–2034. 10.1162/jocn_a_00838

Wan, Q., Menendez, J. A., & Postle, B. R. (2022). Priority-based transformations of stimulus representation in visual working memory. PLOS Computational Biology, 18(6), e1009062. 10.1371/journal.pcbi.1009062

Wang, N., Verschooren, S., Vermeylen, L., Grahek, I., & Pourtois, G. (2025). Hypervigilance strikes a balance between external and internal attention: Behavioral and modeling evidence from the switching attention task. Psychological Research, 89(1), 3. 10.1007/s00426-024-02028-6

Williams, G. C., Nobre, A. C., & Boettcher, S. E. P. (2025). Feature-temporal predictions dynamically modulate performance, feature-based attentional capture, and motor response activity during visual search. Visual Cognition, 33(8), 524–541. 10.1080/13506285.2026.2645176

Xu, Y. (2024). The human posterior parietal cortices orthogonalize the representation of different streams of information concurrently coded in visual working memory. PLOS Biology, 22(11), e3002915. 10.1371/journal.pbio.3002915

Zhou, Y., Curtis, C. E., Sreenivasan, K. K., & Fougnie, D. (2022). Common Neural Mechanisms Control Attention and Working Memory. Journal of Neuroscience, 42(37), 7110–7120. 10.1523/JNEUROSCI.0443-22.2022

